# NADPH phosphatase activity of Mesh1 controls sleep in *Drosophila*

**DOI:** 10.64898/2025.12.14.694245

**Authors:** Taro Ishikawa, Yohei Nitta, Jiro Osaka, Takanari Nemoto, Doshun Ito, Ryosuke Sasaki, Shouta Nonoyama, Akira Oikawa, Atsushi Sugie, Takashi Suzuki, Shinji Masuda

## Abstract

Recent findings have shown that metazoans accumulate the bacterial second messenger guanosine tetraphosphate (ppGpp) and possess *Mesh1*, a gene encoding a ppGpp hydrolase domain. *Mesh1* deficiency affects sleep behavior and eclosion under starvation in *Drosopila*, and ferroptosis in human cells. However, human Mesh1 also exhibits NADPH/NADP^+^ phosphatase activity in vitro, making it unclear whether these phenotypes result from loss of ppGpp hydrolysis, NADPH/NADP^+^ dephosphorylation, or both. To address this, we performed biochemical and genetic analysis of *Drosophila* Mesh1. We first found that *Drosophila* Mesh1 dephosphorylates NADPH, but not NADP^+^, in vitro. We subsequently sought to generate *Drosophila* Mesh1 point mutants specifically impaired in NADPH phosphatase activity. Based on structural data, we mutated W138 and R142, residues that are predicted to interact with NADPH but not ppGpp. W138 was replaced with phenylalanine results in complete loss of NADPH phosphatase activity with retained ppGpp hydrolase activity. Flies carrying the W138F mutation, introduced via genome editing, exhibited shortened total sleep and increased sleep fragmentation in behavioral assays, without changes in intracellular ppGpp levels. These results indicate that the NADPH phosphatase activity of Mesh1, and specifically the W138 residue, is essential for normal sleep regulation in *Drosophila*.

## Introduction

Environmental fluctuations affect physiological functions, behavior, and intracellular metabolism in living organisms. To cope with such changes, organisms have evolved sophisticated regulatory systems that enable adaptive responses to a wide range of environmental conditions. The stringent response is a stress adaptation mechanism conserved in bacteria, and plays a vital role in survival under adverse conditions. Central to this response is the signaling molecule guanosine tetraphosphate (ppGpp), a unique nucleotide that functions as a second messenger ^1^. ppGpp regulates activity of RNA polymerase directly or indirectly, thereby repressing the expression of rRNA, tRNA, and numerous metabolic enzyme genes ^1–5^, while simultaneously promoting the transcription of amino acid biosynthetic genes to compensate for nutrient depletion. These actions collectively reduce energy consumption under stress ^3,6^. Since the stringent response is regulated by intracellular ppGpp levels, bacterial stress adaptation depends on the enzymatic synthesis and hydrolysis of ppGpp. In *Escherichia coli*, these reactions are catalyzed by two enzymes, RelA and SpoT: RelA harbors only a synthetase domain, whereas SpoT contains both synthetase and hydrolase domains ^1,3^. Homologs of these enzymes, collectively known as RelA-SpoT Homologs (RSHs), are conserved across a wide range of organisms, including eukaryotes ^7^.

More recently, genes encoding *Mesh1*, a metazoan homolog of SpoT that retains only the ppGpp hydrolase domain, have been identified in the genomes of several animals, including *Drosophila* and humans ^8,9^. In addition, ppGpp accumulation has been detected in animal cells ^10,11^. However, no enzyme responsible for ppGpp synthesis has been found in animals, and the mechanisms governing ppGpp levels as well as its physiological roles in metazoans remain largely unknown. Intriguingly, a recent study demonstrated that ppGpp levels in the mouse gut microbiota fluctuate in response to the host’s feeding cycle ^12^, suggesting that animals may utilize microbially derived ppGpp even in the absence of intrinsic synthetase enzymes.

In *Drosophila*, deletion of *Mesh1* results in phenotypes such as disrupted sleep behavior and reduced pupation and eclosion rates under starvation conditions ^11^. These phenotypes are accompanied by elevated ppGpp levels in *Mesh1*-deficient flies ^10^, implying that Mesh1 functions to degrade ppGpp in vivo. Moreover, ppGpp accumulation increases under nutrient deprivation, supporting the hypothesis that animals may possess stress response mechanisms involving ppGpp ^11^. Mesh1 knockdown has also been reported to induce the expression of histone deacetylases, leading to repression of *TAZ*, a transcription factor involved in cell cycle regulation ^13^. These findings suggest that Mesh1-mediated cell growth arrest may occur via multiple targets and hint at the presence of a bacterial-like stringent response pathway in metazoans that suppresses cell proliferation under stress ^14^.

In parallel, human Mesh1 (hMesh1) has been shown to exhibit NADPH/NADP^+^ phosphatase activity in vitro, catalyzing the conversion of NADPH to NADH, and NADP^+^ to NAD^+^ ^15^. Due to structural similarities between NADPH and ppGpp, Mesh1 appears capable of hydrolyzing both molecules in vitro. The NADPH/NADP^+^ phosphatase activity of hMesh1 was suggested to regulate ferroptosis by modulating lipid peroxidation through the glutathione system ^14,15^. This enzymatic activity is also required for suppression of *TAZ*-mediated cell proliferation ^13^, raising the possibility that Mesh1 controls intracellular NADPH/NADP^+^ levels, an essential cofactor in numerous metabolic processes.

In this study, we generated a *Drosophila melanogaster* Mesh1 (dMesh1) mutant that selectively lost NADPH phosphatase activity, and investigated how this shift in substrate specificity affects sleep behavior in *Drosophila*. This work not only clarifies the physiological relevance of Mesh1’s dual enzymatic activities, but also provides new insight into the evolutionary conservation of sleep and stress adaptation mechanisms in metazoans.

## Results

### *Drosophila* Mesh1 dephosphorylates NADPH, but not NADP^+^

We first performed biochemical analysis of dMesh1 that was overexpressed in *E. coli* as a his-tagged version (for details, see Materials and Methods). After purification, NADPH/NADP^+^ phosphatase activity of the his-tagged dMesh1 was assessed using the malachite green, which makes a complex with molybdate and released phosphate, whose formation can be monitored by a color change (yellow to green) and absorption increment at 630 nm (Fig. 1a) ^16^. As shown in Fig. 1b, absorption changes at 630 nm could be observed when NADPH used as a substrate. On the other hand, no absorption change was observed with NADP^+^, indicating that dMesh1 specifically dephosphorylates NADPH, but not NADP^+^. Similar observations were reported for hMesh1; it can dephosphorylate both NADPH and NADP^+^, although the activity toward NADP^+^ is approximately 3-fold lower than that toward NADPH ^15^.

**Figure 1.**
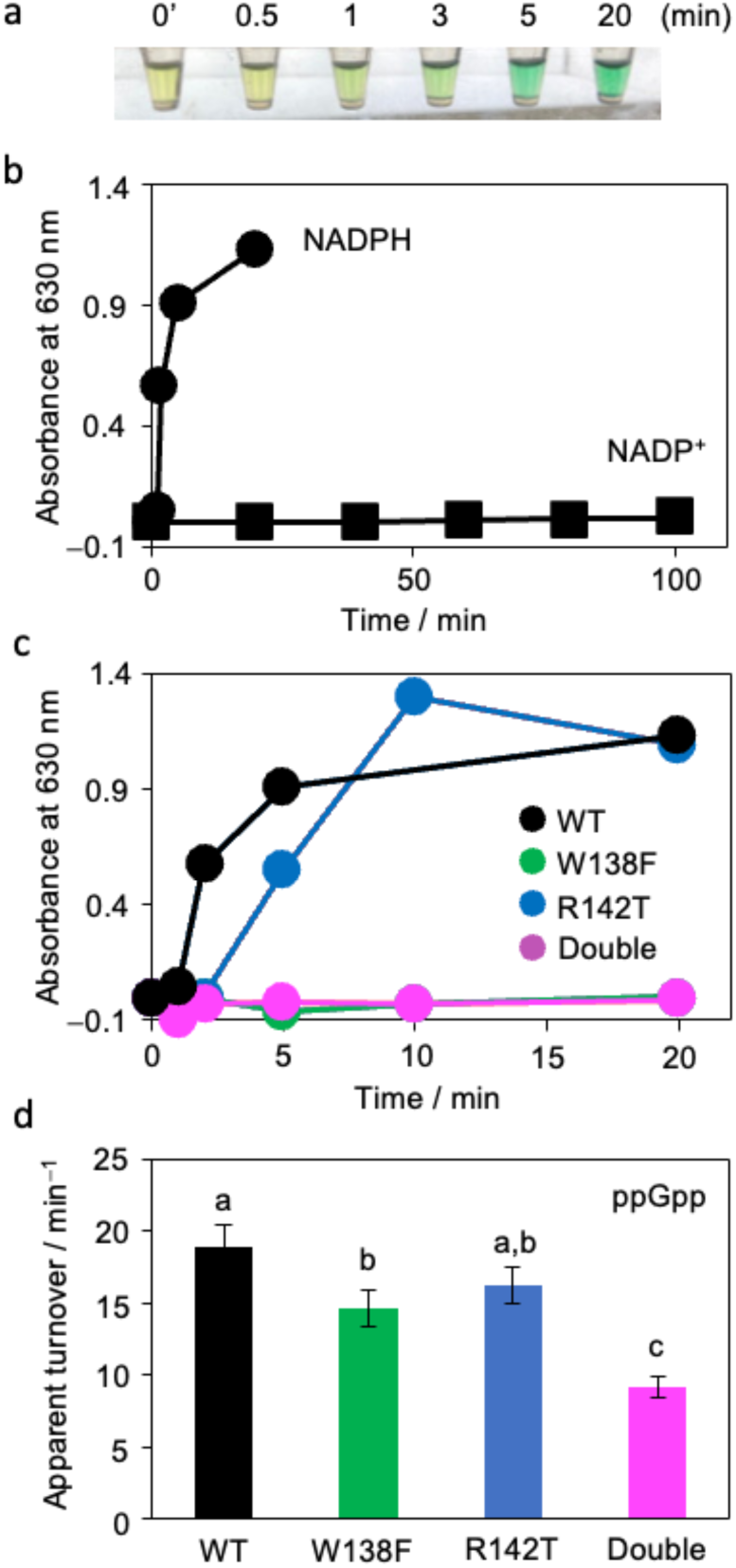
Enzymatic activity of dMesh1. (a) Demonstration of the time course of phosphate release from NADPH catalyzed by dMesh1, detected by a color change (yellow to green) in the malachite green assay. (b) Kinetics of phosphatase activity of WT dMesh1 using 1 mM NADPH or 1 mM NADP⁺ as a substrate. Released phosphate was monitored by the absorbance change at 630 nm due to complex formation with phosphate and malachite green. (c) Kinetics of phosphatase activity of WT, W138F, R142T, and W138F/R142T mutant dMesh1 using 1 mM NADPH as a substrate. Released Pi was monitored as in (b). (d) ppGpp hydrolase activity of WT, W138F, R142T, and W138F-R142T double mutant dMesh1. 1 mM ppGpp was used as a substrate. Data represent means ± SD (n=3). Statistical significance was assessed by Tukey’s test, and groups labeled with identical letters are not significantly different, whereas groups with different letters are significantly different (P < 0.05).

### Designating Mesh1 point mutations affecting NADPH binding

Based on the crystal structure of hMesh1 bound to NADPH (Supplemental Fig. S1) ^15^, we focused on amino acid residues that interact specifically with the nicotinamide moiety of NADPH, a feature absent in ppGpp. Among these residues, W138 and R142 in dMesh1 were found to be conserved exclusively in metazoan Mesh1 proteins, but not in bacterial SpoT homologs (Supplemental Fig. S2 and Supplemental Dataset 1). We hypothesized that point mutations at these positions could disrupt or weaken the interaction with NADPH, and thereby alter Mesh1’s substrate specificity. To this end, we introduced two mutations: W138 was substituted with another aromatic amino-acid phenylalanine (W138F), and R142 was replaced with threonine (R142T), the corresponding residue found in *E. coli* SpoT based on the amino-acid sequence alignment (Supplemental Fig. S2).

We used AlphaFold to predict the structural consequences of these mutations and their potential effects on NADPH binding (Supplemental Fig. S3). The backbone conformations of the variants closely overlapped with that of WT Mesh1, with only minor deviations at the termini. When all four structures were superimposed, the adenosine moiety and the dephosphorylation-target phosphate groups of NADPH were similarly positioned across all variants (Supplemental Fig. S4). In contrast, the orientation of the nicotinamide moiety differed significantly between WT and each of the mutants. These observations suggest that the W138F and R142T mutations alter the mode of NADPH binding to Mesh1, specifically affecting the interaction with the nicotinamide portion.

### Enzymatic activities of Mesh1 mutants

To evaluate the enzymatic activity of dMesh1 mutant variants, three point-mutants (W138F, R142T, and W138F-R142T) together with WT dMesh1 were expressed in *E. coli*, purified, and subjected to activity measurement using the malachite green assay. The R142T mutant retained detectable phosphatase activity toward NADPH, although reaction rate was reduced compared to wild-type Mesh1 (Fig. 1c). In contrast, the W138F single mutant and the W138F-R142T double mutant exhibited no measurable activity, indicating a complete loss of function. These results suggest that the tryptophan at position 138 is essential for NADPH phosphatase activity of dMesh1, while the arginine at position 142 does not appear to be directly involved in catalysis.

We next determined the Michaelis–Menten parameters for NADPH dephosphorylation by wild-type and R142T dMesh1. The *K*_m_ value for NADPH was 0.39 ± 0.04 mM for WT dMesh1 and 2.39 ± 0.60 mM for the R142T mutant. The maximum reaction velocity (*V*_max_) was 2.14 ± 0.07 mM min⁻¹ mg⁻¹ for wild-type and 1.55 ± 0.29 mM min⁻¹ mg⁻¹ for R142T. These data indicate that R142T variant dMesh1 exhibits approximately 6.1-fold lower substrate affinity and 1.4-fold lower catalytic activity toward NADPH compared to the wild-type. The *K*_m_ value for NADPH determined for WT dMesh1 in this study (0.39 ± 0.04 mM) was slightly higher than the previously reported value of hMesh1 (0.11 ± 0.42 mM) ^15^.

We next tested the ppGpp hydrolase activity of WT and various mutant forms of dMesh1 (Fig. 1d). The apparent turnover rate for ppGpp hydrolysis by the W138F mutant was significantly lower than that of WT. A slight reduction in ppGpp hydrolase activity was also observed for the R142T mutant, although the difference compared to WT was not statistically significant. The W138F-R142T double mutant showed a significantly lower ppGpp hydrolase activity than either the WT or the single mutants. These results indicate that the W138F, R142T, and W138F-R142T mutant forms of dMesh1 still retain ppGpp hydrolase activity, although these mutations slightly and negatively affect the activity.

### Phenotypic analysis of *Drosophila* expressing Mesh1 point mutants

Constructs encoding C-terminal FLAG-tagged versions of WT dMesh1 or point mutants (W138F, R142T, and W138F-R142T) were generated and introduced into the endogenous *Mesh1* locus via homologous recombination using CRISPR-based genome editing technique (Fig. 2). To confirm correct insertion, genomic DNA was extracted from the edited *Drosophila* lines and the *Mesh1* locus was PCR-amplified and sequenced. Only flies with confirmed genotypes were used for phenotypic analysis (for details, see Materials and Methods).

**Figure 2.**
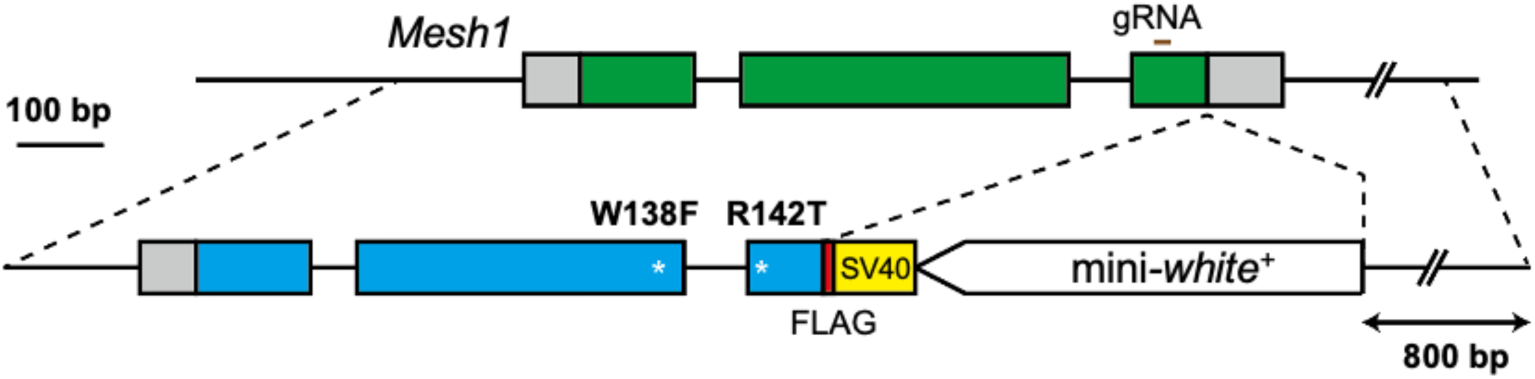
Schematic representation of genome editing for constructing fly lines expressing FLAG-tagged WT dMesh1 and its W138F, R142T, and W138F/R142T mutant variants. The transcription termination signal of SV40 was inserted downstream of the dMesh1 locus. The mini-*white*⁺ gene was used as a marker for homologous recombination. The position of the guide RNA (gRNA) used for double-strand break induction by the CRISPR/Cas9 system is indicated.

At first, we quantified ppGpp levels in *Drosophila* harboring each Mesh1 variant. The measured concentrations were 1.57 ± 0.77 nM mg⁻¹ for WT, 1.21 ± 0.43 nM mg⁻¹ for W138F, 1.46 ± 0.38 nM mg⁻¹ for R142T, and 1.10 ± 0.27 nM mg⁻¹ for the W138F-R142T double mutant (*n*=5). No statistically significant differences were observed among the genotypes. These results indicate, consistent with our initial hypothesis, that the W138F and R142T mutations in dMesh1 do not affect intracellular ppGpp levels in *Drosophila*.

A previous study reported that *dMesh1*-null mutants exhibit reduced sleep levels ^11^. To clarify the physiological functions of Mesh1 that influence sleep, we analyzed the sleep behavior of each point mutant line using the Drosophila Activity Monitoring (DAM) system ^17^. In the DAM system, a single fly is placed in a glass tube with food at one end, and sleep is evaluated by monitoring, using infrared beams, how many times the fly travels back and forth from one end of the tube to the other. For each line, 24–25 individual flies were monitored (for details, see Materials and Methods). After a 3-day acclimation period, sleep behavior was recorded continuously for 3 days.

Figure 3a shows the sleep profiles of each genotype over a 3-day monitoring period. The x-axis indicates time, with alternating white and black bars denoting 12-hour light and dark phases, respectively. The *y*-axis indicates the times of sleep per 60 min. Figure 3b quantifies the sleep ratio separately for the light and dark phases. In WT (Fig. 3a, black line), sleep ratio peaked around ZT8 in the light phase and dropped approaching ZT12, indicating wakefulness at the transition to the dark phase. This was followed by prolonged sleep during the dark phase, continuing until the next light phase, a typical circadian sleep-wake cycle. In contrast, flies carrying the W138F mutation (Fig. 3a, green line) showed similar sleep patterns to WT during the light phase but displayed significantly reduced sleep during the dark phase (Fig. 3b). The R142T mutant (Fig. 3a, blue line) exhibited sleep levels similar to WT during the dark phase but showed significantly increased sleep during the light phase (Fig. 3b). Notably, both W138F and R142T mutants showed a delayed onset of increased sleep at the light-to-dark transition (Fig. 3a). The W138F-R142T double mutant (Fig. 3a, pink line) exhibited significantly reduced sleep in both light and dark phases compared to WT (Fig. 3b).

**Figure 3.**
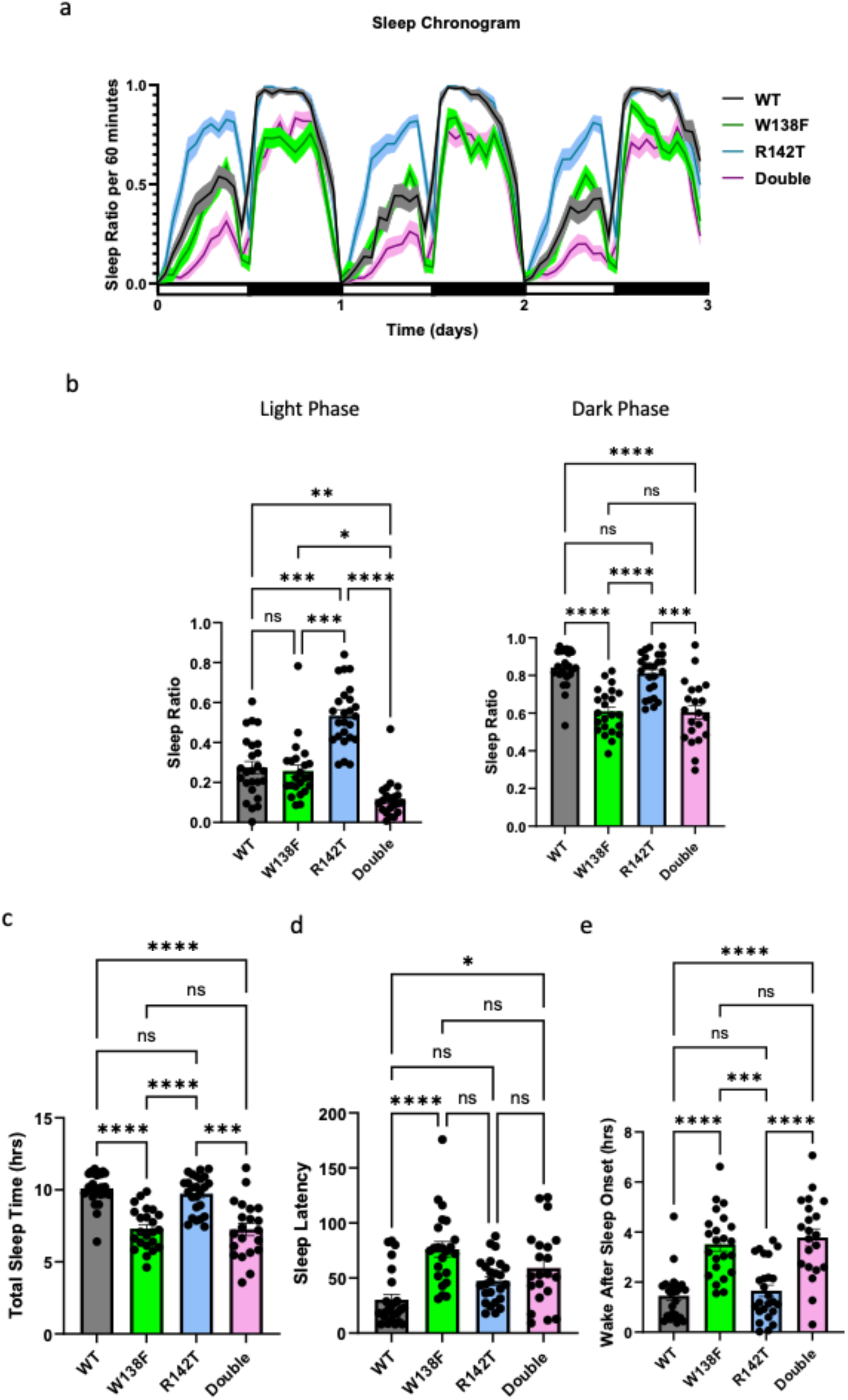
Sleep behavior of WT and mutant flies. (a) Changes in sleep duration per 60 min. (b) Sleep duration per unit time (60 min) during the light and dark phases, respectively. (c) Total sleep duration throughout a 24-h period. (d) Latency to sleep onset after ZT12. (e) Total wake time between the first sleep episode after ZT12 and ZT24. For each line, 24–25 individual flies were analyzed. After a 3-day acclimation period, sleep behavior was recorded continuously for 3 days. ns, no significance. Statistical analyses were performed using the Kruskal-Wallis test with multiple comparisons corrected by Dunn’s method (*, P < 0.05; **, P < 0.01; ***, P < 0.005; ****, P < 0.001; ns, not significant).

Figure 3c compares the total daily sleep time. W138F and the double mutant exhibited a significant reduction in total sleep time compared to WT, whereas R142T mutant were comparable to WT.

Figure 3d shows sleep latency, defined as the time between the onset of the dark phase and the initiation of the first sleep episode. Longer sleep latency indicates greater difficulty initiating sleep. W138F and the double mutant showed significantly increased sleep latency relative to WT, while R142T showed no significant difference. Figure 3e presents the wake after sleep onset (WASO), which reflects the duration of wakefulness between the first sleep episode and the end of the dark phase. Similar to sleep latency, WASO was significantly increased in W138F mutant, but unchanged in R142T.

Taken together, these data demonstrate that both W138F and the W138F-R142T double mutant exhibit consistently reduced sleep duration across both light and dark phases, indicating that the W138 residue is critical for Mesh1-mediated sleep regulation.

### Metabolomic profiling of WT and Mesh1 mutant adults

We next performed metabolome analysis on adult flies of WT and each Mesh1 mutant strain. The results of principal component analysis (PCA) based on the relative abundance of detected metabolites indicated that three principal components (PC1–PC3) accounted for approximately 56% of the total variance (Supplemental Fig. S5). Each genotype (WT, W138F, R142T, or the double mutant) was broadly separated in the PCA plot. These results suggest that the introduction of each mutation affects the overall metabolic composition in a distinct manner.

The number of metabolites that showed a significant increase in abundance relative to WT was 17 for W138F, 1 for R142T, and 2 for the double mutant (Fig. 4a, Supplemental Fig. S6). Conversely, the number of metabolites that were significantly decreased compared to WT was two for W138F, one for R142T, and ten for the double mutant (Fig. 4, Supplemental Fig. S6). Apart from six and two metabolites that were commonly altered in W138F and R142T (either increased or decreased relative to the wild-type), no metabolites exhibited consistent changes across more than one mutant genotype. These findings further support the notion that each Mesh1 mutation differentially impacts the metabolic profile.

**Figure 4.**
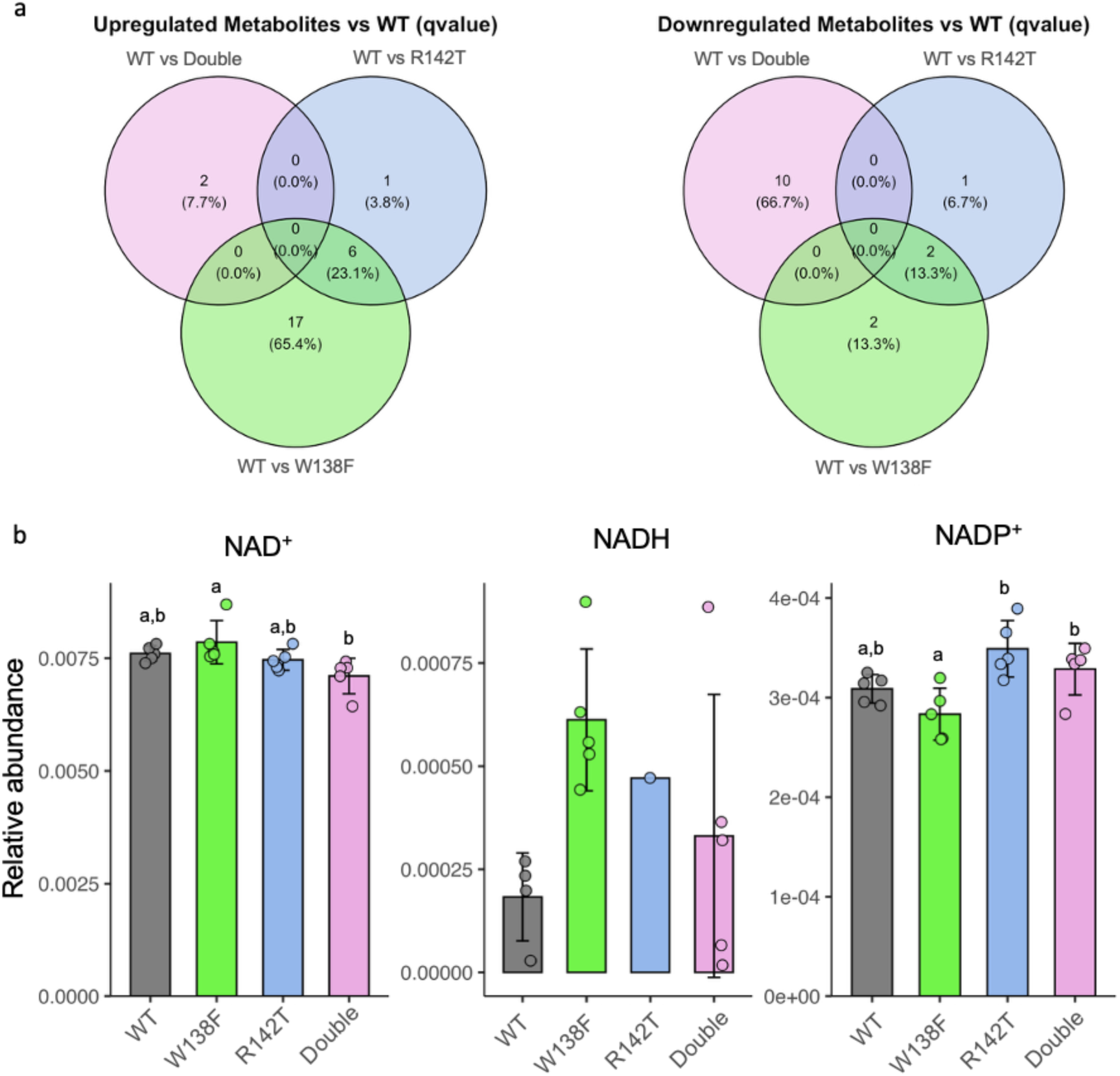
Metabolome analysis of WT and mutant flies. (a) Venn diagrams showing upregulated and downregulated metabolites in each genotype. (b) Levels of NAD⁺, NADH, and NADP⁺ in WT and mutant flies. Statistical significance was assessed by Tukey-Kramer test, and groups labeled with identical letters are not significantly different, whereas groups with different letters are significantly different (P < 0.05). Groups without significant differences are not labeled.

Previous studies have shown that the NADP⁺/NADPH ratio in sleep-promoting neurons plays a critical role in regulating the expression of sleep-related genes, and that alterations in this ratio can modulate sleep behavior ^18^. Based on this, we focused our analysis on the relative levels of NAD⁺, NADH, NADP⁺, and NADPH in the metabolome data. Five biological replicates were analyzed per genotype; however, some samples did not show detectable peak(s) of one or more of these four metabolites. Notably, NADPH was undetectable in all samples by the current metabolome analysis. As shown in Fig. 4b, there were no significant differences in the levels of NAD⁺, NADH, and NADP⁺ between WT and the mutants, although the NAD⁺ level in the W138F mutant was significantly higher than that in the double mutant, and the NADP⁺ levels in the R142T and double mutants were significantly higher than that in the W138F mutant. The NADH level was also elevated in the W138F mutant, although no significance was observed compared to WT. These results suggest that the W138F mutation exerts a slight effect on whole-body NAD⁺, NADH, and NADP⁺ levels in adult flies, which could be at least partially compensated by the R142T mutation.

## Discussion

### Mechanism of Mesh1 phosphatase activity

Biochemical analysis revealed that the tryptophan at position 138, but not the arginine at position 142, is essential for NADPH phosphatase activity (Fig. 1c). These findings provide mechanistic insights by identifying how specific amino acid residues contribute to enzymatic activity. AlphaFold-based structural predictions revealed that none of the mutations (W138F, R142T, or the double mutant) induced major changes in the overall protein structure (Supplemental Fig. S3). Notably, the structural integrity of the region surrounding the NADPH-binding site appeared largely unaffected. Despite these similarities, the W138F and double mutants completely lacked NADPH phosphatase activity, prompting further investigation.

A key difference among the variants was observed in the orientation of the nicotinamide moiety of NADPH (Supplemental Fig. S4, red circles). In all Mesh1 mutants, the nicotinamide group adopted a distinct orientation from that in the wild-type protein, consistent with the fact that the mutations were introduced at residues interacting with this moiety (Supplemental Fig. S1). However, this shift was observed regardless of whether the mutant retained enzymatic activity (e.g., R142T) or not (e.g., W138F), suggesting that structural realignment of the nicotinamide alone does not fully explain the loss of activity. In contrast, the adenosine moiety and the phosphate group targeted for dephosphorylation were similarly positioned across all variants (Supplemental Fig. S4, white circles). This observation implies that the loss of phosphatase activity in W138F and the double mutant may be due to a failure of NADPH to access the substrate-binding site, rather than defects in catalysis itself. Given that the spatial relationship between the active site and the phosphate group was maintained in all predicted structures, NADPH binding to Mesh1 may be sufficient to enable dephosphorylation—provided that proper substrate access is achieved.

Based on these findings, we propose a three-step model for Mesh1-mediated NADPH dephosphorylation: (1) recognition of the nicotinamide moiety by side chains within the α-helical loop, including W138; (2) subsequent access of NADPH to the catalytic site; and (3) dephosphorylation. In the W138F and double mutants, the first step is likely impaired, preventing the downstream reactions from occurring. Comparison of the kinetic parameters between WT dMesh1 and the R142T mutant further supports this model. Notably, the *K*_m_ value was substantially higher in R142T than in WT, while *V*_max_ was only modestly reduced. These results suggest that steps (2) and (3) still occur in R142T, but initial substrate recognition (step 1) is weakened.

A recent report has indicated that Mesh1 binds ppGpp with lower affinity compared to NADPH ^19^, and we here showed that intracellular concentrations of ppGpp in each Mesh1 mutants were the same as those in WT. Taken together, these findings support the view that NADPH phosphatase activity constitutes a primary function of Mesh1 in metazoan metabolic regulation.

### Role of Mesh1 in regulating sleep behavior in *Drosophila*

Flies expressing the W138F or W138F-R142T double mutant form of dMesh1 showed significantly reduced sleep during the dark phase (Fig. 3). These flies also exhibited prolonged sleep latency and increased wake after sleep onset (WASO), phenotypes that closely resemble the sleep deficits previously reported in *Mesh1*-null mutants ^11^. In contrast, the R142T mutant showed sleep parameters largely comparable to those of WT. Combined with our in vitro results showing that the W138F mutation abolishes NADPH phosphatase activity, these behavioral data suggest that the reduction in dark-phase sleep is linked to the loss of NADPH dephosphorylation function in Mesh1.

Given that ppGpp levels remained unchanged across all mutant lines as mentioned in the results section, it is unlikely that the sleep phenotypes observed in *Mesh1*-deficient flies result from loss of ppGpp hydrolase activity. This supports the idea that NADPH, rather than ppGpp, is the primary substrate mediating Mesh1’s physiological functions, at least including sleep regulation, in vivo. Recent findings have demonstrated that the NADP⁺/NADPH ratio in sleep-promoting neurons can influence sleep pressure ^18^. Our results raise the possibility that Mesh1 contributes to sleep regulation by modulating cellular redox status through its control over the NADPH/NADP⁺ and possibly NADH/NAD⁺ balance. Such redox-dependent signaling could impact neuronal activity and sleep pressure at the circuit level.

Metabolome analysis of whole adult flies revealed that only the W138F mutant exhibited elevated, yet insignificantly, NAD^+^ and NADH levels compared with WT (Fig. 4). Given that the W138F mutant dMesh1 lacks NADPH dephosphorylation activity (Fig. 1), this mutation would be expected to result in a decrease, rather than an increase, in NAD^+^/NADH levels. Similarly, the W138F-R142T double mutant also lacks NADPH dephosphorylation activity (Fig. 1) and would be anticipated to exhibit metabolic changes comparable to those observed in the W138F single mutant. However, such changes were not observed, and the double mutant actually restored the phenotype of the W138F mutant (Fig. 4). These findings suggest that the absolute levels and relative ratios of NAD⁺, NADH, NADP⁺, and NADPH are likely influenced by additional regulatory factors, such as de novo synthesis pathways. Furthermore, the effects of dMesh1 point mutations on the levels and redox balance of these pyridine nucleotides may vary across different tissues, including sleep-promoting neurons.

We evaluated sleep behavior during the light phase. Interestingly, R142T mutants displayed significantly increased sleep during this period, while sleep was reduced in the W138F-R142T double mutant (Fig. 3b). Notably, W138F flies exhibited more sleep than the double mutant, suggesting that the R142T mutation is unlikely to contribute directly to increased sleep duration. One possible explanation is that sleep regulation during the light phase may involve a distinct pathway from that of the dark phase, and that Mesh1 participates in both. We also noted a delay in sleep onset at the transition from the light phase to the dark phase in both W138F and R142T mutants, a phenotype that persisted over the 3-day monitoring period. As shown in Fig. 4d, R142T mutants also exhibited a modest increase in sleep latency. However, given that no consistent sleep phenotype was shared exclusively between W138F and R142T across both light and dark phases, the biological significance of this delayed sleep initiation remains unclear.

In conclusion, this study demonstrates that the NADPH phosphatase activity of dMesh1 is essential for normal sleep behavior of *Drosophila*. Sleep is an evolutionarily conserved physiological process, observed from *Caenorhabditis elegans* to humans, and its deprivation is known to cause severe consequences, including cognitive impairments and disruptions in cellular metabolism ^20,21^. Furthermore, sleep disturbances are strongly associated with neuropsychiatric disorders such as bipolar disorder, PTSD, and Alzheimer’s disease, with growing evidence supporting a bidirectional relationship between sleep loss and disease onset ^22–24^. Despite its importance, the mechanisms underlying sleep regulation remain incompletely understood. The neural circuitry of *Drosophila* is considerably simpler than that of humans; however, many fundamental features are conserved, including the use of neuromodulators such as dopamine and serotonin in sleep regulation ^25^. Moreover, *Drosophila* displays circadian-regulated sleep patterns and sleep rebound following deprivation, both of which mirror human sleep behaviors ^26^. Our findings that Mesh1-mediated NADPH dephosphorylation contributes to the regulation of sleep highlight a novel metabolic mechanism underlying sleep homeostasis. These results provide new insights into the biological functions of sleep and may offer a valuable framework for future studies exploring the metabolic regulation of sleep in both invertebrate and vertebrate systems.

## Materials and Methods

### Protein expression and purification

All primers used are listed in Supplemental Table S1. The DNA fragment encoding dMesh1 was synthesized by IDT (USA), which was cloned into pET28a *E. coli* expression vector (Thermo Fisher Scientific). The W138F and/or R142T mutation(s) were introduced by inverse PCR with primer pairs: Mesh1-W138F-F and Mesh1-W138F-R, and Mesh1-R142T-F and Mesh1-R142T-R. These plasmid constructs were separately introduced into *E. coli* expression strain BL21(DE3). The resulting strains were grown at 37°C in LB medium containing 50 mg/L kanamycin. At OD_600_=0.8, 1 mM (final concentration) isopropyl-β-D-thiogalactopyranoside was added to induce protein expression, and further grown for over 16 hours at 16°C. The expressed his-tagged WT and mutant dMesh1 were purified by His-bind region (Thermo Fisher Scientific) according to the manufacturer instructions.

For YjbM expression, DNA fragment encoding *Bacillus subtilis yjbM* was amplified by PCR using primers, YjbM-F-NdeI and YjbM-R-XhoI. The plasmid clone harboring *yjbM* gene whose codons are optimized to express in *Arabidopsis thaliana* ^27^ was used as a template. The amplified DNA fragment was cloned into NdeI-XhoI-cut pET29a (Thermo Fisher Scientific), and the resulting plasmid was introduced into *E. coli* strain BL21(DE3). Expression and purification of his-tagged YjbM is as described above.

### Preparation of ppGpp

Purified 1 µM YjbM was mixed with 5 mM ATP and 5 mM GDP in a buffer containing 100 mM HEPES/NaOH (pH 7.5), 200 mM NaCl, 20 mM MgCl_2_ and 20 mM KCl, and incubated for 16 h at 37°C. YjbM was precipitated by the addition of one volume of chloroform. The aqueous phase that contained the nucleotides was diluted with double-volume of distilled water and was subjected to anion exchange chromatography (Hi-trap Q; GE Healthcare). A gradient ranging from 0-2 M NaCl was used for elution of the nucleotides. ppGpp eluted at around 200 mM NaCl was further purified by solid-phase extraction protocol as described previously ^28^.

### Enzymatic analysis of dMesh1

NADPH/NADP^+^ phosphatase activity of dMesh1 was monitored by the malachite green assay ^16^. For the assay, the malachite green solution was prepared as follows. At first, 60 ml of concentrated sulfuric acid was mixed with 300 ml of deionized water, followed by addition of 0.44 g of malachite green, and the resulting solution was named as dye solution. On the day of use, 2.5 ml of 7.5% (w/v) ammonium molybdate was added to 10 ml of dye solution followed by 0.2 ml of Tween 20. The resulting solution was designated as the malachite green solution. All NADPH/NADP^+^ phosphatase activity measurements were carried out in a total volume of 800 µl in a buffer containing 100 mM Tris/HCl (pH 7.5), 200 mM NaCl, 20 mM MgCl_2_ and 20 mM KCl. For each assay, 1.25 µM dMesh1 and varying concentration of NADPH or NADP^+^ (as indicated in figure legends) were used. All assays were carried out at 30°C. At each time point, 160 µl of reaction solution was mixed with 40 µl the malachite green solution. After incubated for 10 min, absorption at 630 nm was measured, and the concentration of Pi was estimated based on the calibration curve obtained with varying concentration of KH_2_PO_4_. Enzyme parameters were calculated by determining the initial reaction rates of each Mesh1 variant at NADPH concentrations of 0.2, 0.4, or 0.8 mM. The *V_max_* and *K_m_* values were calculated by Enzyme Kinetic Calculator ^29^.

ppGpp hydrolase activity of dMesh1 was monitored by HPLC. All reactions were carried out in a total volume of 40 µl of a buffer containing 100 mM Tris/HCl (pH 7.5), 200 mM NaCl, 20 mM MgCl_2_ and 20 mM KCl. For each assay, 1.25 µM dMesh1 and 1 mM ppGpp were used. Assays were carried out at 30°C. At each time point, 40 µl of reaction solution was mixed with 10 µl of 0.5 mM EDTA to stop the reaction. HPLC measurements were performed with an Elite LaChrom Series system (Hitachi) and a 5 µm SAX column (SphereClone 150 X 4.6 mm; Phenomenex). After running 20 min with a buffer containing 50 mM NH_4_H_2_PO_4_, a gradient up to 100% of 500 mM NH_4_H_2_PO_4_ over 20 min was applied. The reaction product GDP and the remaining substrate ppGpp were detected at a wavelength of 254 nm, in agreement with standards.

### Generation of genome editing vectors and targeting constructs

All primers used are listed in Supplemental Table S1. The FLAG epitope tag as well as W138F, R142T and W138F-R142T mutations were introduced into the *Mesh1* locus by the genome editing technique established previously ^30^. We first modified pHT009 carrying *white* to generate plasmid constructs introducing FLAG-tag as well as Mesh1 point mutations (Fig. 2). DNA fragments for FLAG-tagged Mesh1 WT and W138F, R142T and W138F-R142T mutants were synthesized by IDT. DNA fragments encoding 0.8 kbp *Mesh1* N-terminal region and SV40 transcription termination sequence were amplified by PCR using primer pairs; Mesh1-up-F-HindIII and Mesh1-up-R, and FLAG-SV40-term-F and SV40-term-F. The two PCR-amplified fragments were mixed with synthesized DNA encoding FLAG-tagged WT Mesh1, those of W138F and W138F-R142T mutants, and then cloned into HindIII restriction site of pHT009 using In-Fusion coning regent (Clontech, USA). The Mesh1 C-terminal UTR region was amplified by PCR using the primer pair, Mesh1-dw-F-KpnI and Mesh1-dw-R-KpnI, which was then coned into the KpnI site (upstream of *white*) of the plasmids constructed above, and the resulting constructs were named pHT009-Mesh1-WT, pHT009-Mesh1-W138F and pHT009-Mesh1-double.

The plasmid for guide RNA synthesis was constructed below. The primers, delta-Mesh1-F and delta-Mesh1-R were mixed, incubated at 98°C for 5 min, and then cooled down to 37°C by 1 hour. The annealed DNA was cloned into BbsI-cut pBFv-U6.2 ^30^, and the resulting plasmid was named pBFv-mesh1.

pHT009-Mesh1-WT, pHT009-Mesh1-W138F and pHT009-Mesh1-double were separately injected with pBFv-mesh1 into CS, *w*^1118^ (*w^−^*). Transformants were produced by BestGene Inc. (U.S.A). Flies were kept in standard *Drosophila* medium at 25 °C. Candidate strains inducing homologous recombination at the *Mesh1* locus were screened by the presence and absence of the *white* maker. Insertion of FLAG-tag as well as W138F, R142T and W138F-R142T mutations in *Mesh1* locus were confirmed by sequencing of the PCR products amplified with the genomic DNA of the candidate strains. We isolated R142T single mutant line from strains produced by injection of pHT009-Mesh1-double and pBFv-mesh1.

### Fly Husbandry

Flies were reared on a standard cornmeal–yeast medium containing 23 g agar, 94 g granulated sugar, 190 g glucose, 101 g yeast, 230 g cornmeal, 2.25 g calcium chloride, and 27.15 g potassium bitartrate per 2 L water. The mixture was heated at 90 °C for 15 min, and after cooling to 60 °C, 6.015 g nipagin dissolved in 30 mL ethanol was added. The medium was dispensed into vials to a depth of approximately 2 cm and allowed to solidify.

Genome-edited lines were maintained as balanced stocks, but homozygous flies were selected based on phenotypic markers and used for the analyses.

### Physiological analysis of *Drosophila*

Sleep behavior of *Drosophila* strains were performed with the DMA system (TriKinetics). Each homozygous male fly from of different genotypes was picked and inserted into a glass tube (ϕ 5 mm X 65 mm) in which food was placed at one end. An artificial circadian rhythm was established by alternating 12-hour light and dark phases. Time was indicated using Zeitgeber Time (ZT), with the onset of the light phase designated as ZT0. Sleep was defined as a period of inactivity lasting 5 minutes or longer (i.e., no infrared beam crossings), and death was defined as continuous inactivity for more than 12 hours. Data analysis was performed using Rtivity ^30^.

### Metabolite analysis

Quantification of ppGpp in adult *Drosophila* strains were performed with by ultra-performance liquid chromatography coupled with a tandem quadruple mass spectrometer equipped with electrospray interface as described previously ^10,11^. Metabolome analysis of *Drosophila* strains was performed by capillary electrophoresis time-of-flight mass spectrometry as described previously ^31^.

Metabolite intensities obtained from metabolomic analysis were normalized to the fresh weight of each *Drosophila* sample to calculate the relative intensity per unit fresh weight. For statistical comparison, normalized intensities were compared between WT and experimental groups. Metabolites showing more than a twofold increase or less than a half-fold decrease relative to WT, with a Benjamini–Hochberg (BH) adjusted *p*-value (qvalue) < 0.05, were defined as differentially expressed metabolites (DEMs) ^32^. Venn diagrams were subsequently generated based on the identified DEMs.

### Statistics and reproducibility

Statistical significance of data was tested by Tukey’s, Tukey-Kramer or Kruskal-Wallis tests with multiple comparisons corrected by Dunn’s method test by R. The sample size (*n*) and the nature of replicates have been given wherever relevant.

## Acknowledgment

We thank the Integrative Bioscience Facility at Institute of Science Tokyo for technical assistance. We also thank Noriko Nomura for her technical support.

