## Supplemental Tables and Figures for "NADPH phosphatase activity of Mesh1 controls sleep in *Drosophila*"

**Supplemental Table S1**. Primers used in this study (5’->3’)

| | |
|---|---|
| delta-Mesh1-F | CTTCGTCGTGATCAGTACTTCGTT |
| delta-Mesh1-R | AAACAACGAAGTACTGATCACGAC |
| Mesh1-dw-F-KpnI | GTTCTCGAGCGACTGGTACCACGATCAAAAA CCAACAAATTACAG |
| Mesh1-dw-R-KpnI | CTATAGGGCGAATTGGTACCACGACAGGAGA CCTGGAAGCTGG |
| Mesh1-up-F-HindIII | CGAATTGGCCAAGCTTGGCGGAGGAGGCCG CCAAAGCAGCG |
| Mesh1-up-R | CCCTGTCGGTGTGTTGACTTGCAGA |
| SV40-term-F | GATTATAAAGATGACGATGACAAATAAGGATC TTTGTGAAGGAACCTTA |
| SV40-term-R | TGGGTCTAGTGGATCCAGACATGAT |
| Mesh1-W138F-F | GGGATTTACTCAGGAACGTCGCGATCAGTAC |
| Mesh1-W138F-R | TCCTGAGTAAATCCCGTTGGTGTGTTGACCTG |
| Mesh1-R142T-F | CAGGAAACTCGCGATCAGTACTTCGTGTGGG |
| Mesh1-R142T-R | ATCGCGAGTTTCCTGAGTCCATCCCGTTGGTG |
| White-F | GAATACATTTAATTTAGAAAATGCTTGG |
| Mesh1-up-F2 | TTTGATTGGCATTTCCTACTACATC |

**Supplemental Table S2**. Accession numbers of Mesh1 homologs used for constructing the amino-acid sequence alignment shown in Supplemental Fig. S2

| Organism | Accession No. |
|---|---|
| Homo sapiens | NP_001273380 |
| Mus musculus | Q9D114.1 |
| Drosophila melanogaster | Q9VAM9.1 |
| Xenopus tropicalis | Q28C98.1 |
| Danio rerio | Q568P1.1 |
| Poeciliopsis prolifica | JAO16771.1 |
| Ceratitis capitata | JAB92171.1 |
| Labeo rohita | RXN38077.1 |
| Camelus dromedarius | KAB1257087.1 |
| Hypsibius dujardini | OQV25566.1 |
| Heterocephalus glaber | JAO00752.1 |
| Strongyloides ratti | XP_024506477.1 |
| Thamnophis elegans | XP_032088609.1 |
| Lingula anatina | XP_013406804.1 |
| Gigaspora margarita | KAF0473293.1 |
| Nitrosarchaeum | GBL42451.1 |
| Bacillus subtilis | WP_019715221.1 |
| Brucella melitensis | ALM34099.1 |
| Burkholderia multivorans | AJY15143.1 |
| Caulobacter vibrioides | WP_010919427.1 |
| Chlamydia trachomatis | CPR68030.1 |
| Corynebacterium glutamicum rel | CAF20036.1 |
| Corynebacterium glutamicum relH | CAF20012.1 |
| Dictyoglomus thermophilum | ACI19252.1 |
| Enterococcus faecalis | WP_033640712.1 |
| Gloeobacter violaceus | BAC90689.1 |
| Meiothermus silvanus | WP_013156925.1 |
| Oceanithermus profundus | WP_013457017.1 |
| Pseudomonas aeruginosa | VVH82182.1 |
| Rhodobacter capsulatus | WP_013069013.1 |
| Thermoanaerobacter ethanolicus | EGD52207.1 |
| Trichormus variabilis ATCC 29413 | ABA24586.1 |
| Xylella fastidiosa | WP_038227877.1 |

| | |
|---|---|
| Arabidopsis thaliana RSH2 | NP_188021.1 |
| Arabidopsis thaliana RSH3 | NP_564652.2 |
| Arabidopsis thaliana CRSH | NP_188374.2 |

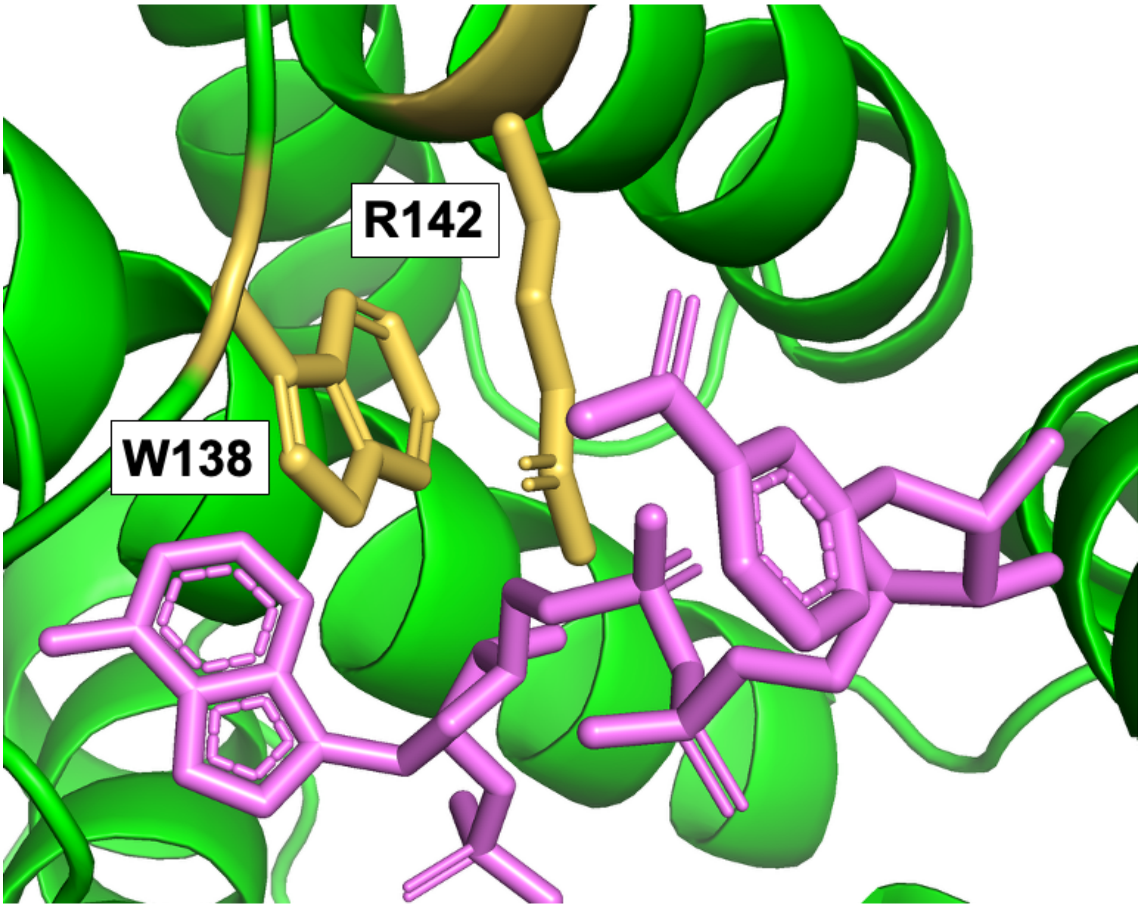


**Supplemental Figure S1.** Structure of the hMesh1-NADPH complex (PDB: 5VXA). The architecture of the apo-protein is shown in the ribbon diagram. NADPH and side chains of W138 and R142 are shown in the stick model (colored in purple and dark yellow, respectively).

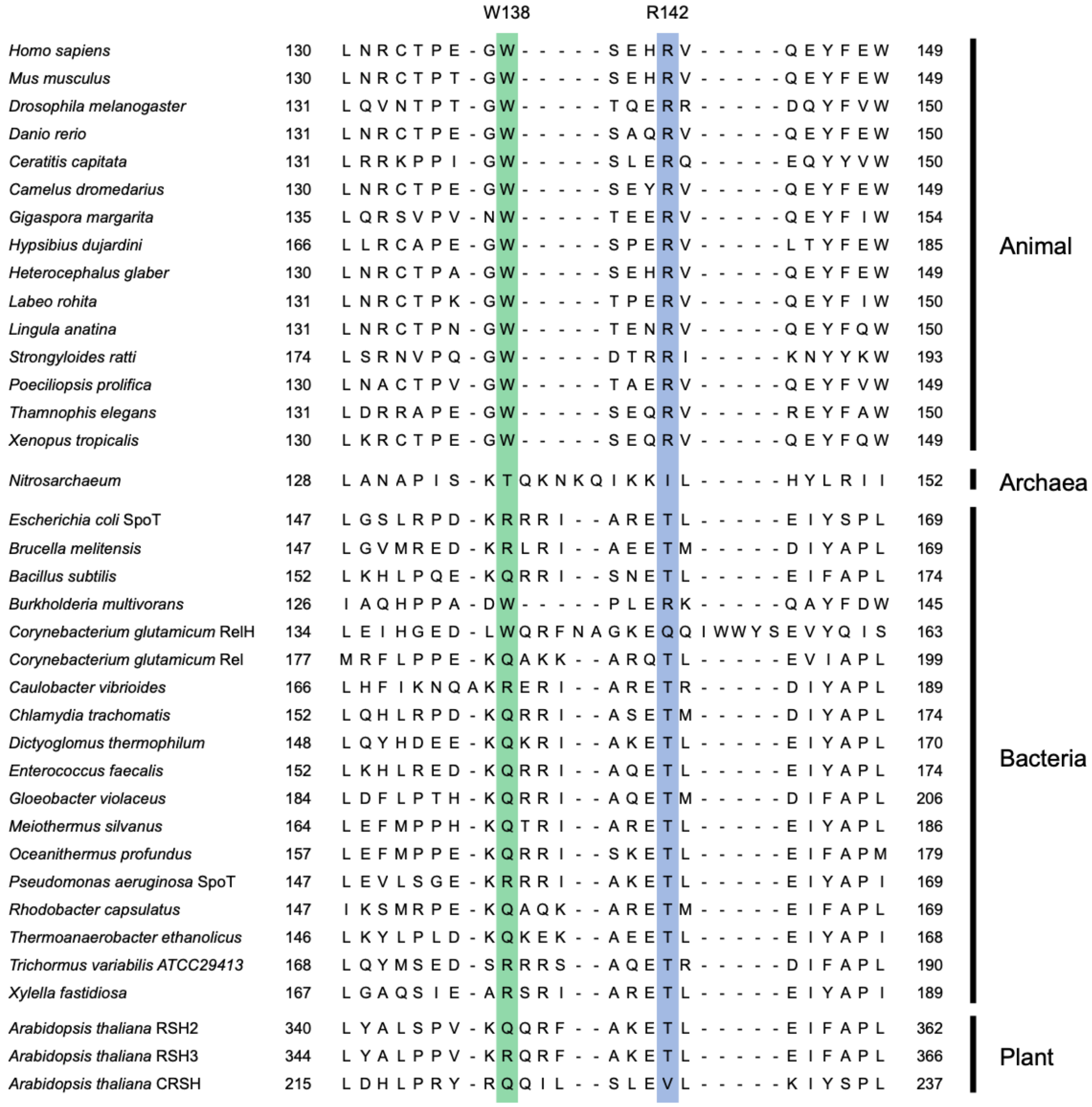


**Supplemental Figure S2.** A partial amino acid sequence alignment of Mesh1 and bacterial SpoT homologs. Accession numbers of the sequences were shown in Supplemental Table S2. The full-length alignment is shown in Supplemental Dataset 1.

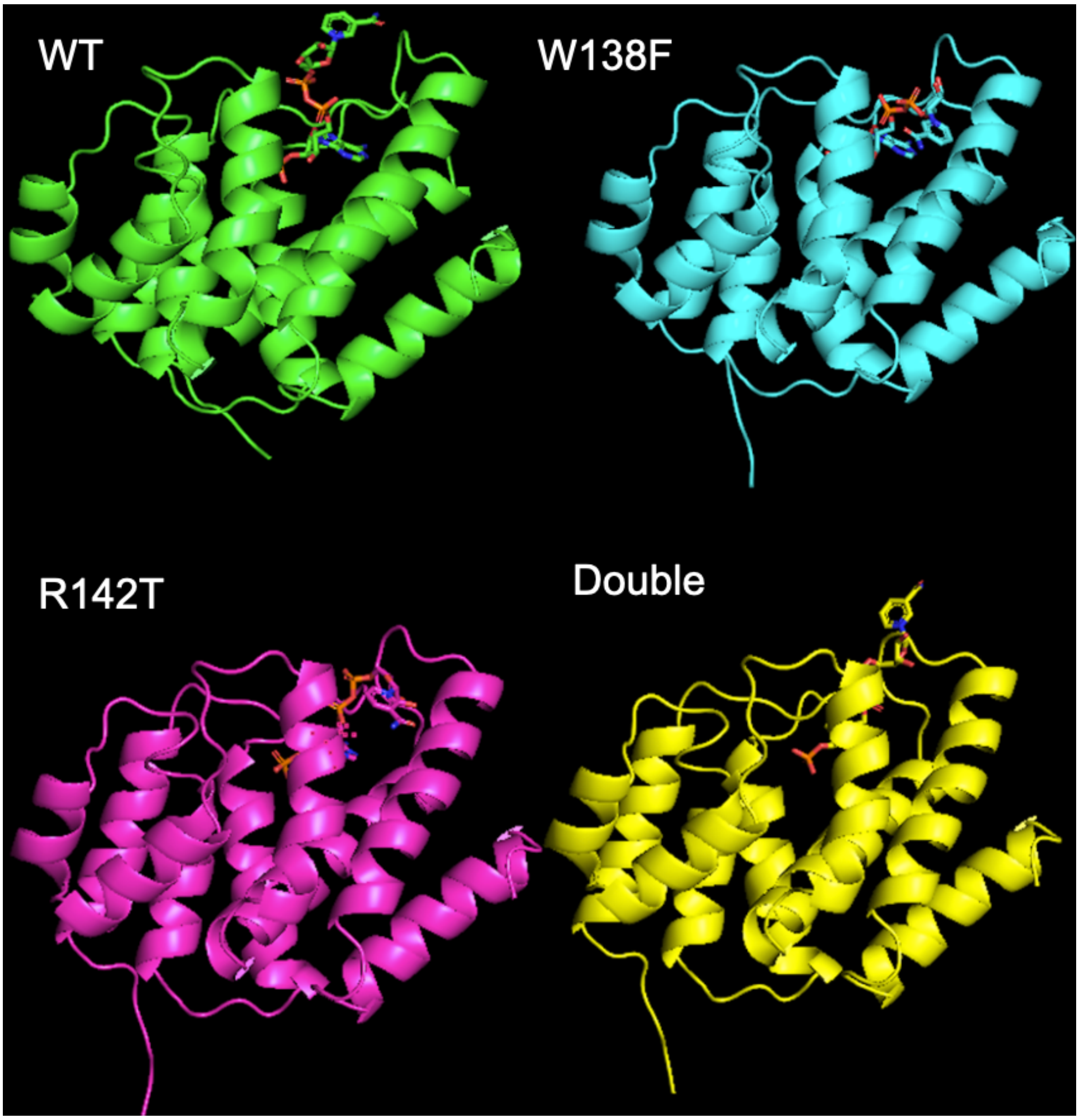


**Supplemental Figure S3.** Predicted structures of the dMesh1-NADPH complex estimated by AlphaFold2. The architecture of the apo-protein is shown in the ribbon diagram. NADPH and side chains of W138 and R142 are shown in the stick model.

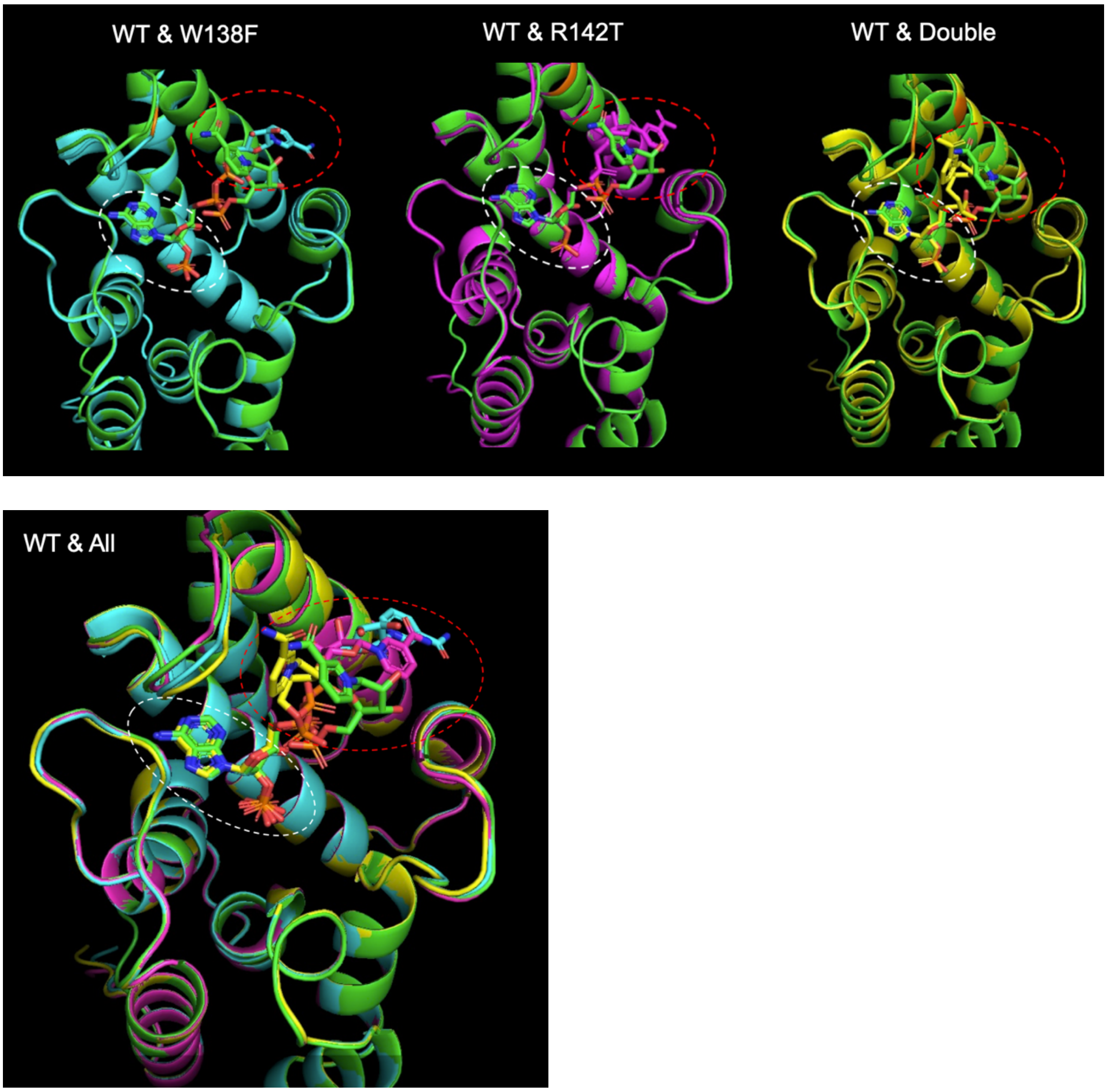


**Supplemental Figure S4.** Superimposed structures of the predicted dMesh1–NADPH complexes estimated by AlphaFold2. The architectures of the apo forms are shown as a ribbon diagram, and NADPH and the side chains of W138 and R142 are represented as stick models. The apo forms of WT, W138F, R142T, and W138F-R142T double mutant are colored in green, cyan, purple, and yellow, respectively. NADPH molecules bound to each variant are shown in the same respective colors. Positions of the adenosine diphosphate moiety and the nicotinamide moiety of NADPH are indicated by white-dotted and red-dotted circles, respectively.

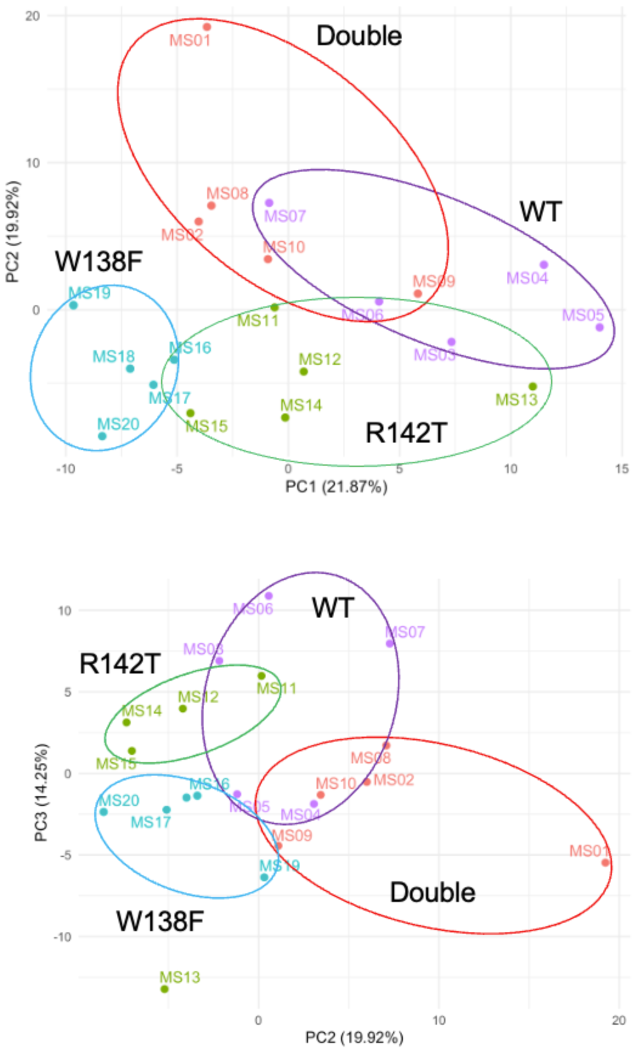


**Supplemental Figure S5.** Principal component analysis of metabolites in adult flies of *Drosophila* WT and W138F, R142T and W138F-R142T double mutants. Experiments were repeated five times, and the data from each experiment were plotted separately.

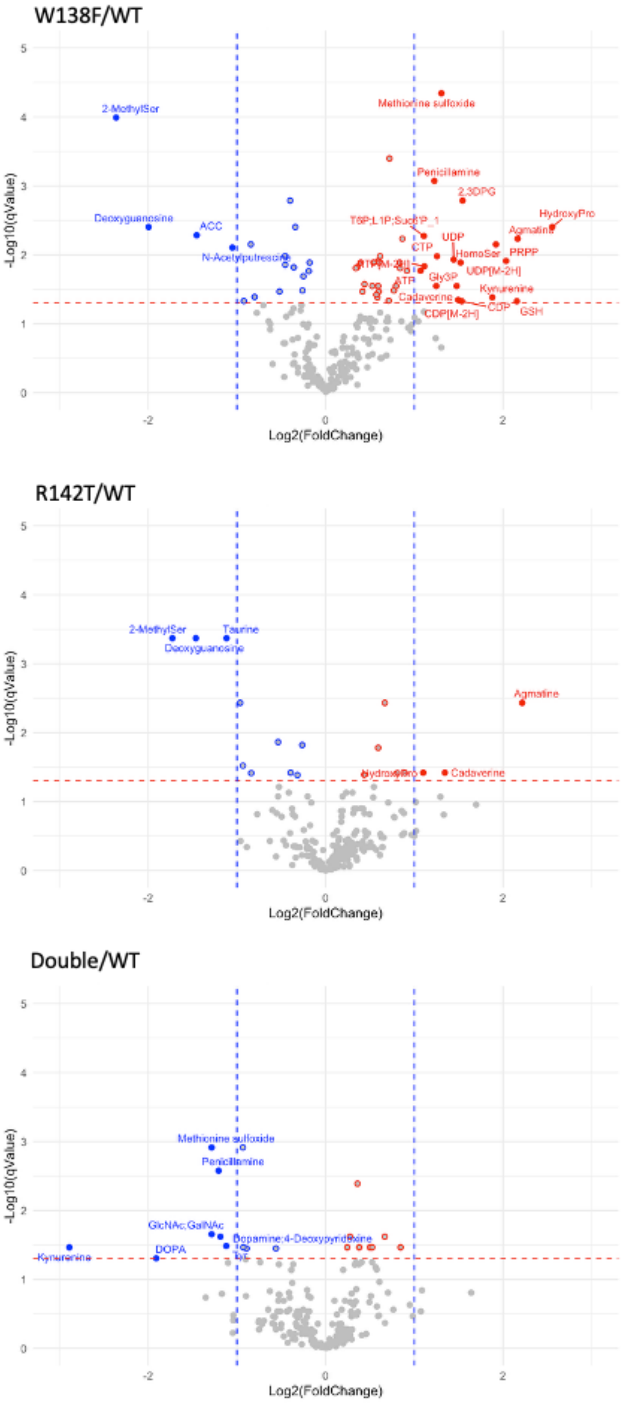


**Supplemental Figure S6.** Volcano plots comparing metabolites in adult flies of *Drosophila* WT and W138F, R142T or W138F-R142T double mutants. Metabolites whose levels were significantly downregulated and upregulated compared to those in WT are indicated in blue and red, respectively.

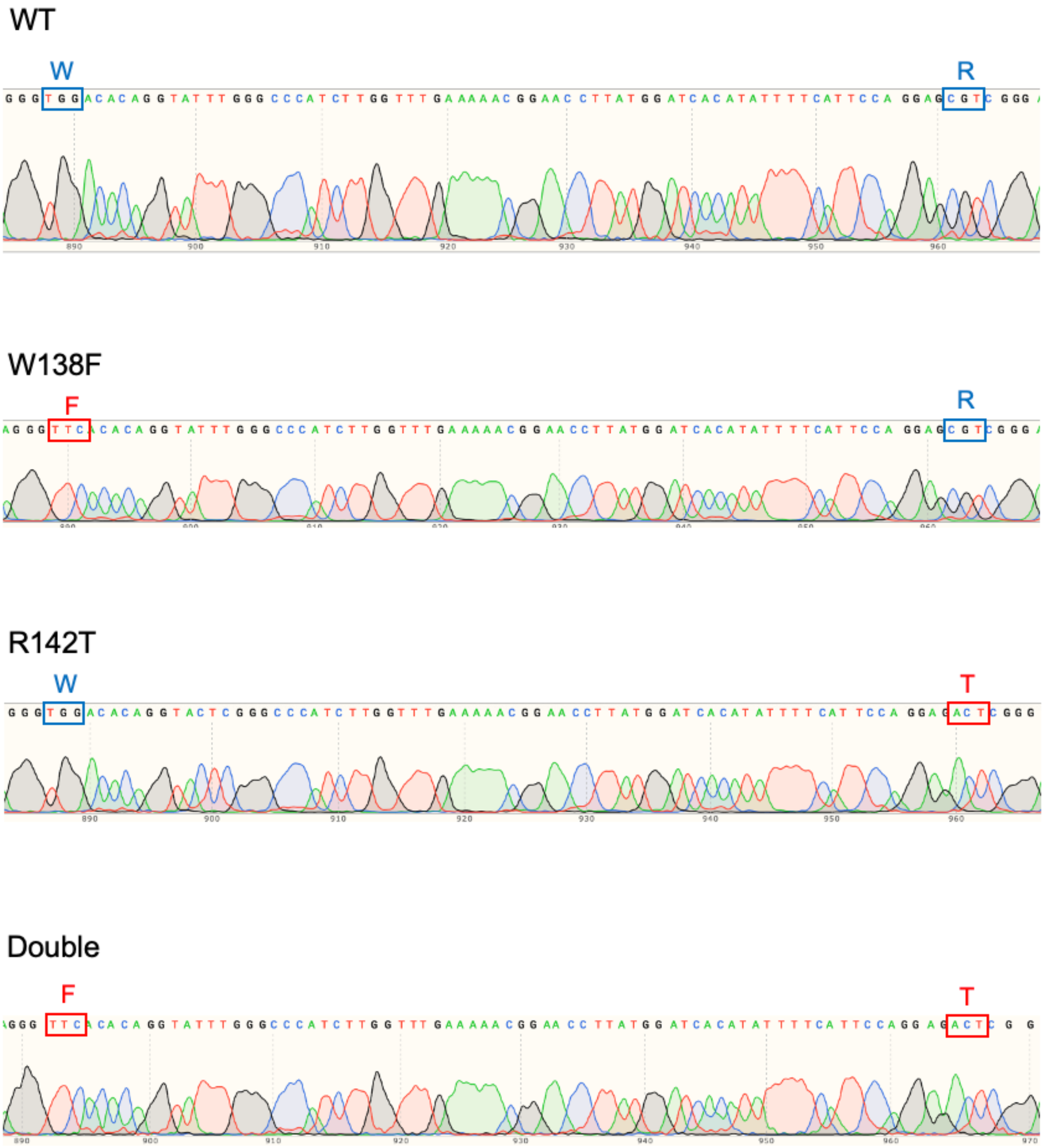


**Supplemental Figure S7.** Sequencing results confirming genotypes of W138F, R142T and W138F-R142T double mutations. Mesh1-coding regions were PCR amplified with genomic DNAs isolated from homozygous flies of each genotype and the primer pair, white-F and Mesh1-up-F2 (Supplemental Table S1). The amplified fragments were directly sequenced by the primer Mesh1-up-F2.
